# Nitrogenase inhibition limited oxygenation of the Proterozoic atmosphere

**DOI:** 10.1101/475236

**Authors:** John F. Allen, Brenda Thake, William F. Martin

## Abstract

Cyanobacteria produced the atmospheric O_2_ that began accumulating 2.4 billion years ago^1^, leading to Earth’s Great Oxidation Event (GOE)^2^. For nearly 2 billion years following the GOE, O_2_ production was restricted and atmospheric oxygen remained low^2–5^. Oxygen rose again sharply with the advent of land plants roughly 450 million years ago, which increased atmospheric O_2_ through carbon burial^4–5^. Why did the O_2_ content of the atmosphere remain constant and low for more than a billion years despite the existence of O_2_-producing cyanobacteria? While geological limitations have been explored^2–7^, the limiting factor may have been biological, and enzymatic. Here we propose that O_2_ was kept low by oxygen inhibition of nitrogenase activity. Nitrogenase is the sole N_2_-fixing enzyme on Earth, and is inactive in air containing 2% or more O_2_ by volume^8^. No O_2_-resistant nitrogenase enzyme is known^9–12^. We further propose that nitrogenase inhibition by O_2_ kept atmospheric O_2_ low until upright terrestrial plants physically separated O_2_ production in aerial photosynthetic tissues from N_2_ fixation in soil, liberating nitrogenase from inhibition by atmospheric O_2_.

---

Current views of oxygen in Earth history (Fig. 1) depict the first traces of O_2_ appearing in the atmosphere starting about 2.7 to 2.5 Gy ago^1–5^. During the Great Oxidation Event, or GOE, roughly 2.4 billion years ago^2^, O_2_ rose to about 10% of its present atmospheric level (PAL), corresponding to an atmosphere of roughly 2% O_2_ by volume^2^, or even less^3^. Isotopic studies indicate that for roughly 1.5 billion years following the comparatively sudden GOE, further net O_2_ accumulation ceased, with atmospheric levels remaining stable and below 10% PAL^2–4^.

**Fig. 1.**
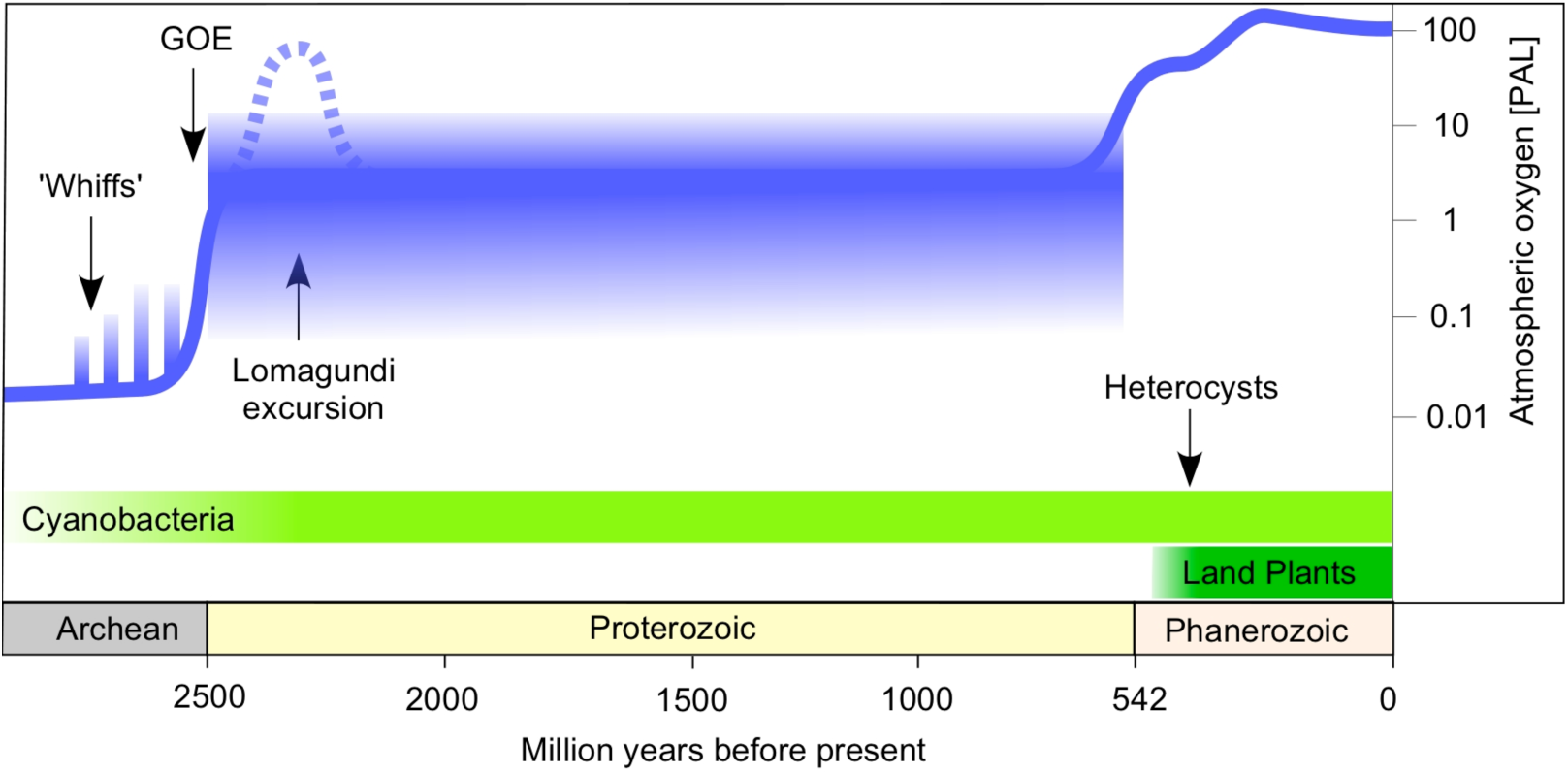
Schematic summary of O_2_ accumulation in Earth history. Modified after refs. 1-5. For most of the Proterozoic eon, free O_2_ was much less abundant than it is today. Lyons et al. (2014) estimate Proterozoic O_2_ in the atmosphere as low as 0.001 PAL while Holland^2^ estimates atmospheric O_2_ at around 0.1 to 0.2 PAL. Stolper and Keller^4^ estimate mid-Proterozoic deep ocean dissolved O_2_ concentrations at about 11 µM or roughly 0.06 of the present value of 178 µM. “Whiffs” refers to isotope signatures for evidence of transient, local O_2_ before the GOE^1,3^. The Lomagundi excursion is represented as a dotted line because it is included in the summary of ref.^3^ but not in that of refs.^1,2,4,5^. Heterocysts are differentiated cells of some cyanobacteria, and protect nitrogenase from inactivation by O_2_. Their relevance is that cyanobacteria have an ancient fossil record, but the oldest fossil heterocysts^26^ are younger than land plants, suggesting that cyanobacteria evolved this mechanism of O_2_ protection in response to Phanerozoic O_2_ accumulation. PAL; Present Atmospheric Level. GOE; Great Oxidation Event.

With atmospheric O_2_ low, marine O_2_ stayed low as well. Geochemical evidence suggests that the oceans remained largely anoxic throughout the Proterozoic^1–5^, with a rapid rise to roughly modern oxygen levels starting around 580 My ago, perhaps as recently as only 430 My ago^4,5^ (Fig. 1). Late increases in atmospheric O_2_ implicate the emergence of land plants and terrestrial carbon burial as a causal factor. Today, land plants comprise roughly 97% of Earth's surface-exposed biomass^7^. Their ecological success has been linked to the rise in O_2_ because terrestrial sequestration of organic carbon as fibrous biomass protects it from reoxidation to CO_2_, curbing O_2_ consumption^4–5^.

What limited oxygen accumulation? A major puzzle of O_2_ history is why O_2_ rose so late, that is, why atmospheric and marine O_2_ levels stayed low for almost 2 billion years despite the existence of cyanobacteria, which were capable of continuous light-driven O_2_ production. What held cyanobacteria back, why did O_2_ stop accumulating after the GOE and why did it remain low during the Proterozoic, or the “Boring Billion” as it is sometimes called^3^.

Geochemists have long recognized that Proterozoic O_2_ stasis presents a problem and have proposed a number of explanations to account for the delayed oxygen rise. Some proposals posit a steady supply of geochemical reductants from within the Earth, such as Fe^2+^ or S^2–^, reductants that continuously consumed the O_2_ produced by cyanobacteria, keeping O_2_ low^8^. Other proposals invoke biotically induced changes affecting in the degree of mixing between nutrient rich reservoirs and the photic zone, for example through animal burrowing activity^9^. Germane to many proposals is the concept that crucial nutrients such as molybdenum, which is required for nitrogenase activity, were limited in supply by geochemical factors and that nitrogenase limited primary production for this reason^6^. These proposals and others^2–5^ might apply to some areas of the ocean or some phases of Earth’s history. But how and why any set of factors should act in concert to keep O_2_ low for almost 2 billion years is yet unresolved^10^.

Nitrogenase regulates oxygen levels. We propose that O_2_-dependent feedback inhibition of a single enzymatic activity limited O_2_ accumulation during the boring billion: inhibition of nitrogenase by O_2_ gas. Carbon and nitrogen enter the biosphere in distinct chemical reactions catalysed by specific enzymes. For carbon there are six pathways of CO_2_ assimilation that differ in age, oxygen tolerance, and key CO_2_ reducing enzymes^11^. For N_2_ there is only one entry point into metabolism: nitrogenase^12–14^. Nitrogenase is widespread among cyanobacteria^12^. There are Mo, Fe and V containing isoforms of the enzyme that all share a common ancestor and homologous active sites^13–15^.

The nitrogenase active site is replete with metal cofactors (Fig. 2) and harbours a metal coordinated carbide carbon atom, unique among all enzymes known so far^14^. Like a blacksmith, nitrogenase uses ancient but robust technology. Nitrognease has an obligatory H_2_ producing side reaction, and it requires 8 electrons and 16 ATP per N_2_ fixed, the ATP being consumed at steps that alter the redox potential of FeS clusters via conformational change^13^. Nitrogenase requires numerous assembly factors^14^, and has been neither replaced nor improved during evolution, which reveals that the solution that life found to fix N_2_ is the only one readily attainable in 4 billion years of physiological engineering by microorganisms. Nitrogenase is a limiting factor. It is inhibited by O_2_ in a feedback loop (Fig. 3), and this simple property alone could limit O_2_ accumulation over geological time.

**Fig. 2.**
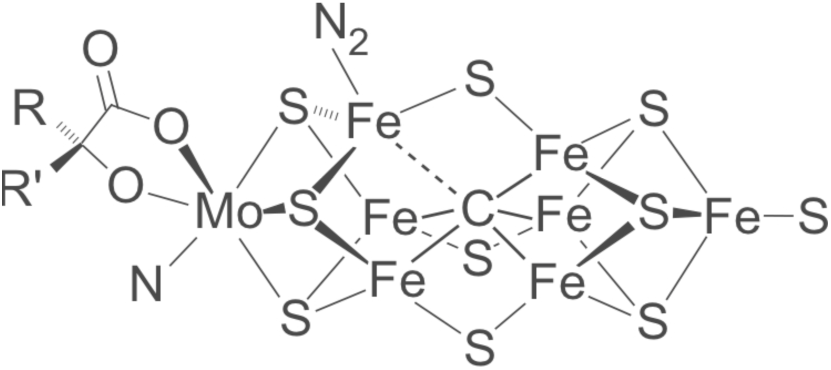
Model of the oxygen-sensitive active site of nitrogenase. Redrawn from ref.^15^ with the proposed binding site for N_2_.

**Fig. 3.**
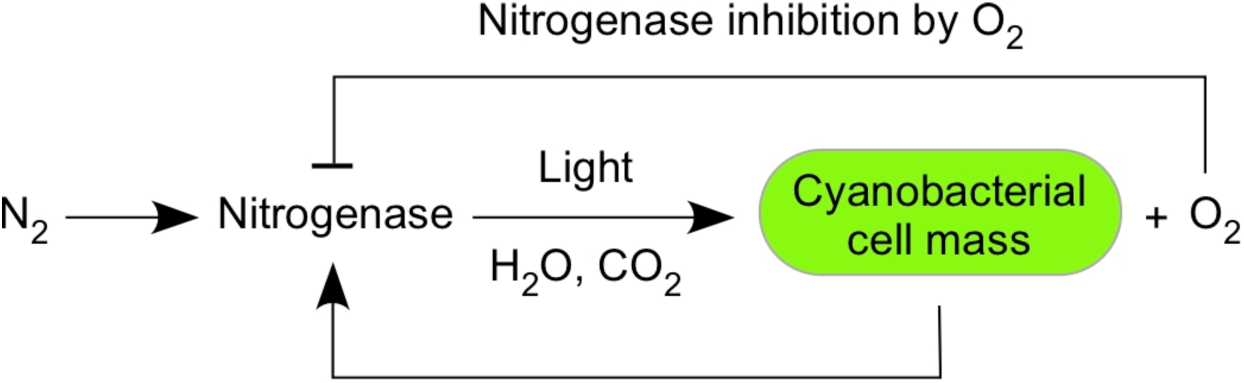
Inhibitory feedback at nitrogenase. O_2_ inhibits nitrogenase, which is required for O_2_ production in photosynthesis. A steady state is reached at environmental O_2_ levels not exceeding 10% PAL.

Nitrogenase feedback inhibition operates as follows. By dry weight, cells are about 50% carbon and about 10% nitrogen. Cyanobacteria had water as an unlimited reductant for CO_2_fixation, but, for net growth to occur, N_2_ incorporation had to keep pace. Nitrogenase is inhibited by oxygen, the product of water oxidation — but there is a threshold of oxygen concentration below which nitrogenase remains active and above which nitrogen fixation ceases completely. If diazotrophic cyanobacteria are grown under conditions where they have sufficient CO_2_ and light, and with N_2_ as the sole N source, then they grow and accumulate no more than 2% oxygen in their culture atmosphere^16^. The 2% O_2_ remains constant during prolonged culture growth because this is the O_2_ partial pressure beyond which nitrogenase activity becomes inhibited. With greater O_2_, nitrogenase is inactivated and there is no fixed N to support further biomass accumulation. With less O_2_, nitrogenase outpaces CO_2_ fixation until the latter catches up, returning O_2_ to 2% in the culture. In cyanobacteria, CO_2_ fixation means O_2_ production.

In microbial mats, oxygen inhibits nitrogenase activity and nitrogenase gene transcription following the onset of illumination during the natural diel cycle^17^. The initial effect of illumination, however, is to increase nitrogenase activity to its maximum value by means of increased ATP and reductant from photosynthetic electron transport. After a lag of a few hours, O_2_ concentration becomes inhibitory to nitrogen fixation^17^. Because there is no biochemical alternative to nitrogenase for fixing N_2_, because there are no O_2_ tolerant nitrogenases known, and because reductant for CO_2_ fixation was not limiting for cyanobacteria, this feedback loop would have operated, on a planetary scale, for two billion years or more. While primary production using H_2_S instead of H_2_O as in *Oscillatoria limnetica*^18^ is also subject to O_2_-feedback inhibition, it would have been limited via the availability of reductant at O_2_ levels far below those created by of oxygenic photosynthesis ^19^, and would not have impacted O_2_ accumulation. Nitrogenase is an O_2_ inhibited sensor that kept environmental O_2_ low throughout the Proterozoic.

Cyanobacteria have evolved mechanisms to avoid nitrogenase inhibition by oxygen^12^, including N_2_ fixation in the dark^20^, heterocysts^21^ or filament bundles as in *Trichodesmium*^22^. Critics might counter that any one of those mechanisms could have bypassed O_2_ feedback inhibition. There are three problems with this objection. First, evolution operates without foresight. Second, the mechanisms that cyanobacteria use to deal with modern O_2_ levels appear to have arisen independently in diverse phylogenetic lineages, not at the base of cyanobacterial evolution when water oxidation had just been discovered^23,24^. Third, the oldest uncontroversial fossil heterocysts trace to land ecosystems of the Rhynie chert and are merely Devonian in age^25^ (Fig. 1), suggesting that heterocysts arose late in evolution, probably in response to levels of O_2_ exceeding 2% by volume. Fossil akinetes — cyanobacterial resting spores — have been found in older sediments^26,27^, yet there is no direct evidence for heterocysts older than the first land plants.

The concept of limiting metal availability (Mo, V, or Fe) for nitrogenase activity^6^ is an element of many proposals to account for low Proterozoic O_2_. Our proposal differs from nutrient limitation in a crucial aspect. Limiting the number of active nitrogenase enzymes in the environment by limiting metal (Mo, V, and Fe) availability only limits the rate at which cyanobacteria produce O_2_, requiring other factors to impose limits upon the final O_2_ partial pressure. Nitrogenase feedback inhibition regulates the O_2_ partial pressure directly, independently of the rate of photosynthesis, and generates a value that corresponds to the geochemical observation.

An O_2_ overshoot 2.3 billion years ago is suggested by an isotopic anomaly called the Lomagundi excursion. At 2.3 to 2.2 Ga ago, the isotopic record first reported from the Lomagundi formation in Zimbabwe indicates burial of heavy (^13^C enriched) carbon^3^. This ^13^C increase is interpreted^3^, though not universally^2^, as indicating the presence of large amounts of O_2_ on a global scale. If that interpretation is correct, its least explicable aspect is that following the Lomagundi excursion, oxygen levels drop once again^3^. Yet they do not drop to pre-cyanobacterial levels, rather they drop to oxygen levels very near 2% O_2_, the oxygen partial pressure that nitrogenase feedback inhibition generates. If the Lomagundi excursion is taken as a valid proxy for high global O_2_ levels, the following situation prevailed at the GOE. O_2_ is a strong oxidant. Its contribution to metabolic evolution was not just new metabolic pathways, but more complete oxidation of existing organic substrates^28^. O_2_ mobilized organic nitrogen and carbon that had been sequestered in biomass. By liberating sequestered nitrogen (and carbon as CO_2_) that had previously been inaccessible to anaerobes, the onset of O_2_ accumulation at the GOE initiated a positive growth feedback loop for aerobic autotrophs that were not reductant limited: cyanobacteria. When anaerobically deposited nitrogen reserves had been liberated, nitrogenase feedback inhibition set in, driving O_2_ levels down to Proterozoic levels, and keeping them low for a billion years thereafter. Our proposal does not hinge upon the Lomagundi excursion, yet if the excursion is interpreted as evidence for transiently high global O_2_ levels then our proposal can account both for its emergence(nitrogenase independent N availability during the excursion) and for its decline in the subsequent return to low O_2_.

When and why did feedback inhibition at nitrogenase cease to keep O_2_ low? At the origin of land plants, the nature of biomass changed and O_2_ production by upright terrestrial plants became physically separated from N_2_ fixation in aquatic environments and soil. Deposition by land plants of nitrogen-depleted cellulose, billions of tonnes of it, became a massive sink for CO_2_ without exerting similar effects on nitrogen availability, thus allowing O_2_ to increase through the standard mechanism of carbon burial, bypassing control by aquatic nitrogenase feedback.

Critics might interject that O_2_ levels began to rise before the first fossil occurrence of land plants^29^. We point out that nitrogenase limitation determines the maximum O_2_ partial pressure near the water surface for nitrogenase-limited oxygen production. This limit does not identify the timepoint at which the deep ocean becomes fully oxic, since that depends upon other factors such as reductant load, ocean mixing, or both, independently of photic zone nitrogenase limitation. Stolper and Keller^4^ report that deep ocean oxygenation became complete 540 million years ago. If so, that was the first time (or possibly the first time since the Lomagundi excursion) that N-rich organic ocean floor sediment came into widespread contact with oxygenated water. This contact released organic N, leading to atmospheric O_2_ increase, after which O_2_ levels dropped once again^29,30^ to the value imposed by the nitrogenase limit. Nitrogenase inhibition returns O_2_ to low levels following O_2_ increases, thus explaining an otherwise puzzling aspect of Proterozoic O_2_ variation.

In conclusion, oxygen inhibition of any ecosystem’s cornerstone enzyme activity, nitrogenase, created a bottleneck for oxygenic primary production that is sufficient to account for low oxygen levels throughout the boring billion. Nitrogenase feedback inhibition could directly account for Proterozoic low oxygen stasis. It would have driven down transiently higher O_2_ levels ensuing from nitrogenase-independent N availability, and it would have ceased at the origin of land plants. Our model requires light, CO_2_ and N_2_ in the photic zone and hence accommodates local and global variation in geochemical conditions while remaining robust to their effects. We propose that the factor limiting Proterozoic O_2_ accumulation was not geochemical. It was biological, and the attribute of a single enzyme, nitrogenase, contained within and synthesized by living cells.

## Acknowledgements

We thank Olivia P. Judson and Dan Wang for comments on the manuscript. JFA thanks the Leverhulme Trust for Research Fellowship EM-2015-068. WFM thanks the ERC (grant no. 666053), the Volkswagen Foundation (grant no. 93 046) and the DFG (grant no. 1426/21-1) for funding.

